# ScaleSurfer: multi-scale anatomical segmentation and parcellation of the human brain

**DOI:** 10.64898/2026.07.01.735927

**Authors:** Ryan Hammonds, Cindy Chen, Bradley Voytek

## Abstract

Human brain magnetic resonance imaging (MRI) revolutionized our ability to non-invasively probe individual differences in neuroanatomy. These anatomical scans, in turn, also allow us to accurately localize functional MRI (fMRI) activity. However, extracting anatomical labels and structural characteristics, such as cortical surface area or thickness, is a computationally demanding task, taking on the order of hours per brain volume. This is an intrinsically multi-scale problem given that local image structure defines fine boundaries, whereas accurate assignments depend on broader anatomical context. Here, we introduce ScaleSurfer, a three-dimensional convolutional vision transformer model based on multi-scale learning. Convolution blocks capture local anatomical detail and a transformer bottleneck integrates the distributed spatial context. This approach provides rapid, whole-brain morphometric feature estimation, including volume, cortical thickness, surface area, and curvature. Importantly, ScaleSurfer accomplishes this nearly five orders of magnitude faster than current pipelines, taking 150-500 ms instead of 5 hours. We validated ScaleSurfer on multiple datasets, showing stable learning across heterogeneous MRI collections, and demonstrate feasibility by training an interpretable Alzheimer’s disease classifier that identifies reductions in primarily medial temporal lobe subregions compared to healthy controls. ScaleSurfer positions multi-scale representation learning as a practical route toward faster, anatomically faithful structural MRI processing, whose speed paves the way for nearly real-time anatomical quality control during scanning.

## 1 Main

### 1.1 Segmentation & Parcellation

Structural magnetic resonance imaging (MRI) is foundational for the non-invasive study of the human brain. As technology improves, we can now identify incredibly fine-scale differences in brain morphology at the sub-millimeter scale [1, 2]. Neuroanatomical differences are apparent across neurological [3, 4], psychiatric [5, 6], and neurodevelopmental disorders [7–9]. In addition to their diagnostic utility, high resolution structural scans permit us to better localize neural activity using functional MRI (fMRI) and their associated co-active networks [10–12]. This localization is critical for studying the neural basis of human behavior and cognition. As foundational as the analysis of structural MRI is, it is computationally intensive and depends on preprocessing pipelines that transform anatomical regions into labeled regions and regional morphometric estimates.

The most widely used method for segmentation, cortical surface reconstruction, and parcellation is FreeSurfer [13–17]. Related ecosystems in SPM, FSL, and ANTs rely on similar approaches for automated structural MRI preprocessing [18–20]. However, FreeSurfer takes several hours to process neuroanatomical scans. Deep learning has the potential to reduce these massive computational requirements and processing times. Structural MRI segmentation methods such as DeepNAT [21], QuickNAT [22], AssemblyNet [23], FastSurfer [24, 25], SynthSeg [26], and deepmriprep [27] have demonstrated that large parts of anatomical preprocessing can be replaced by end-to-end model inference, and perform comparably to established pipelines.

While these approaches have substantially reduced the computational cost of anatomical preprocessing, they are primarily designed to accelerate or approximate existing segmentation workflows. Here, we introduce ScaleSurfer, a three-dimensional convolutional vision transformer for rapid multi-scale brain morphometry. In our approach, convolutional blocks capture local anatomical structure and a transformer bottleneck integrates distributed spatial context to estimate region-wise volume, cortical thickness, surface area, and curvature. This is most similar to 3D-TransUNet [28], but applied to a FreeSurfer segmentation setting. UNesT [29] is also conceptually related, using a hierarchical transformer encoder to achieve segmentation. ScaleSurfer builds upon segmentation models and directly predicts subject-level regional morphometric summaries derived from cortical parcellation and subcortical segmentation. These features include regional volume, cortical thickness, surface area, number of vertices, and curvature and folding measures, which are widely used in group analysis, biomarker discovery, and predictive modeling [30–32].

While prior deep learning models directly predicted region volume and thickness measures [33], ScaleSurfer leverages multi-scale learning to predict the full FreeSurfer set of volumetric, surface, and global features, with processing times on the order of hundreds of milliseconds per brain. We show that ScaleSurfer-derived morphometric features are highly correlated with FreeSurfer estimates that are derived from hours of processing time per brain – nearly five orders of magnitude slower than ScaleSurfer. Finally, we show that ScaleSurfer can be used to extract interpretable neuroanatomical features in a classification task, distinguishing patients with Alzheimer’s disease from healthy control participants.

### 1.2 Multi-Scale Learning

Whole-brain anatomical segmentation is intrinsically multi-scale. Local contrast, tissue boundaries, and fine-scale morphology determine where gross neuroanatomical structures begin and end, but correct labeling depends on the broader anatomical context of the whole brain, such as which hemisphere, the relative position of the lobes, midline (a)symmetries, local relationships, and so on. That is, similar features can correspond to different anatomical regions depending on the broader context, such as in the medial temporal lobe where the amygdala, hippocampus, entorhinal cortex, and parahippocampal gyrus can have similar local contrast, but are distinguished by their position within the larger anterior-posterior and medial-lateral organization of the temporal lobe.

Here, we leverage the fact that convolutional encoder-decoder models excel at multi-scale learning. Such models provide a useful inductive bias for anatomical segmentation, because their architecture assumes that images are locally structured, that features can be reused across space [34–38], which is particularly useful for neuroanatomical parcellation and segmentation. Convolution alone only acquires long-range structure indirectly through depth, pooling, and receptive-field growth, whereas transformers provide a mechanism for explicitly modeling long-range interactions [39–41]. In medical image segmentation, this has led to a broad family of hybrid multiscale models, including TransBTS [42], CoTr [43], UNETR [44], Swin UNETR [45], nnFormer [46], UNesT [29], MISSFormer [47], TransUNet [48] and 3D-TransUNet [28].

### 1.3 ScaleSurfer

ScaleSurfer (Fig. 1) produces FreeSurfer parcellation and segmentation labels using multi-scale learning. The network follows the logic of 3D-TransUNet-like models where a 3D convolutional encoder learns local multi-scale features, such as the fine anatomical boundary between tissue types. Then, a transformer bottleneck aggregates broader context over a compressed latent token embeddings, learning at the global scale, between distant regions. A decoder then restores voxel-wise spatial precision [28, 44, 48], integrating representations across various scales. The resulting ScaleSurfer output is a voxel-wise prediction of 118 segmentation and parcellation labels.

**Fig. 1.**
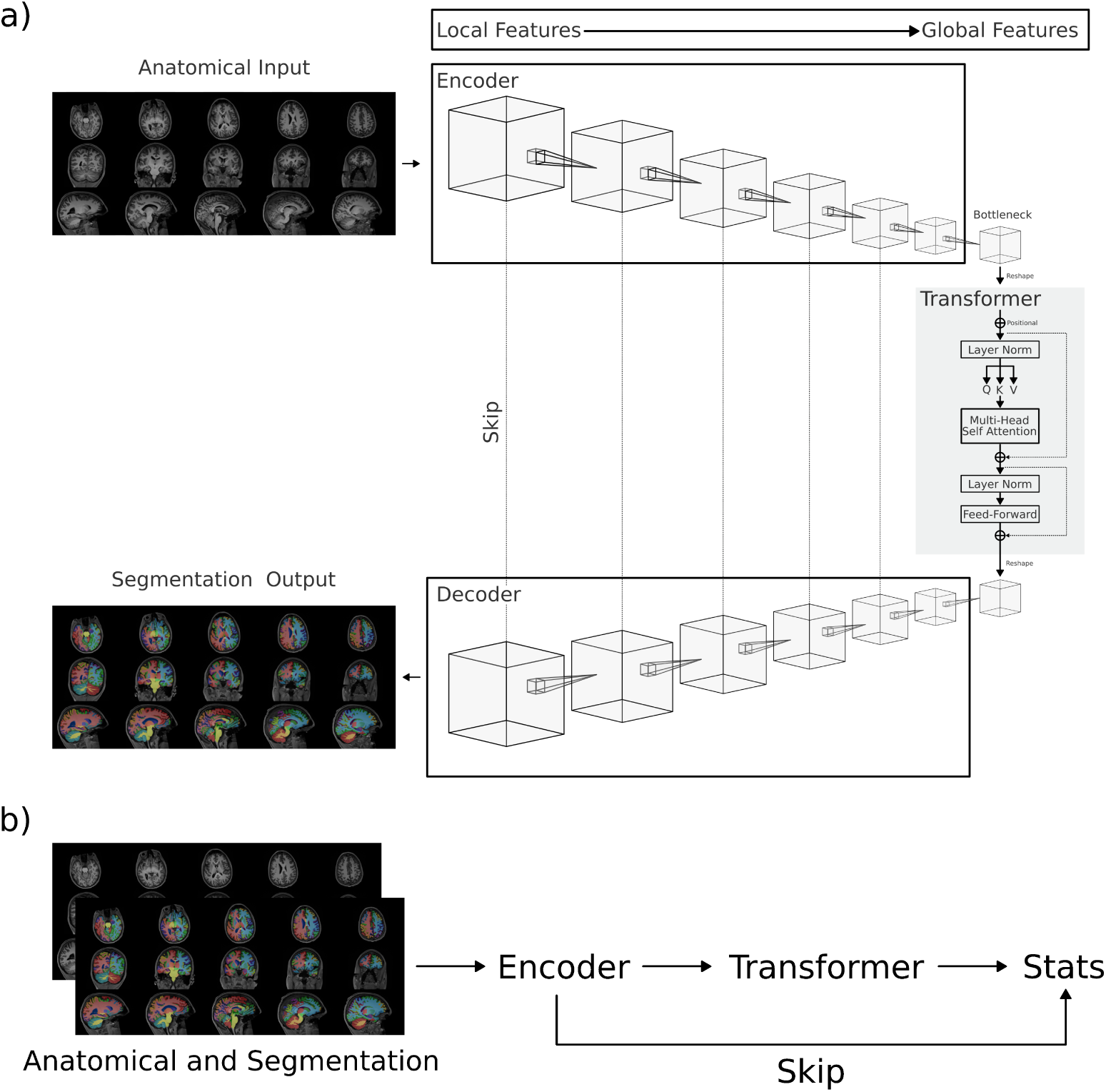
ScaleSurfer. **a)** A 3D convolution encoder learns local features. Each subsequent convolution layer learns increasingly global features as the receptive field grows. The transformer bottleneck learns global features. Each visual token has a large receptive field and attends to every other visual token, facilitating global learning. Skip connections pass multi-scale feature representations from the encoder to the decoder. The decoder is trained to classify each voxel with a segmentation or parcellation label. **b)** Anatomical images and predicted labels are then used to predict region-wise statistics: volume, thickness, surface area, number of vertices, curvature, and the folding index. Global measures, e.g, intracranial volume, are also predicted. The same encoder and transformer architecture are used as in the segmentation model, while the voxel-wise decoder is replaced with model heads for each morphometric feature type.

**Fig. 2.**
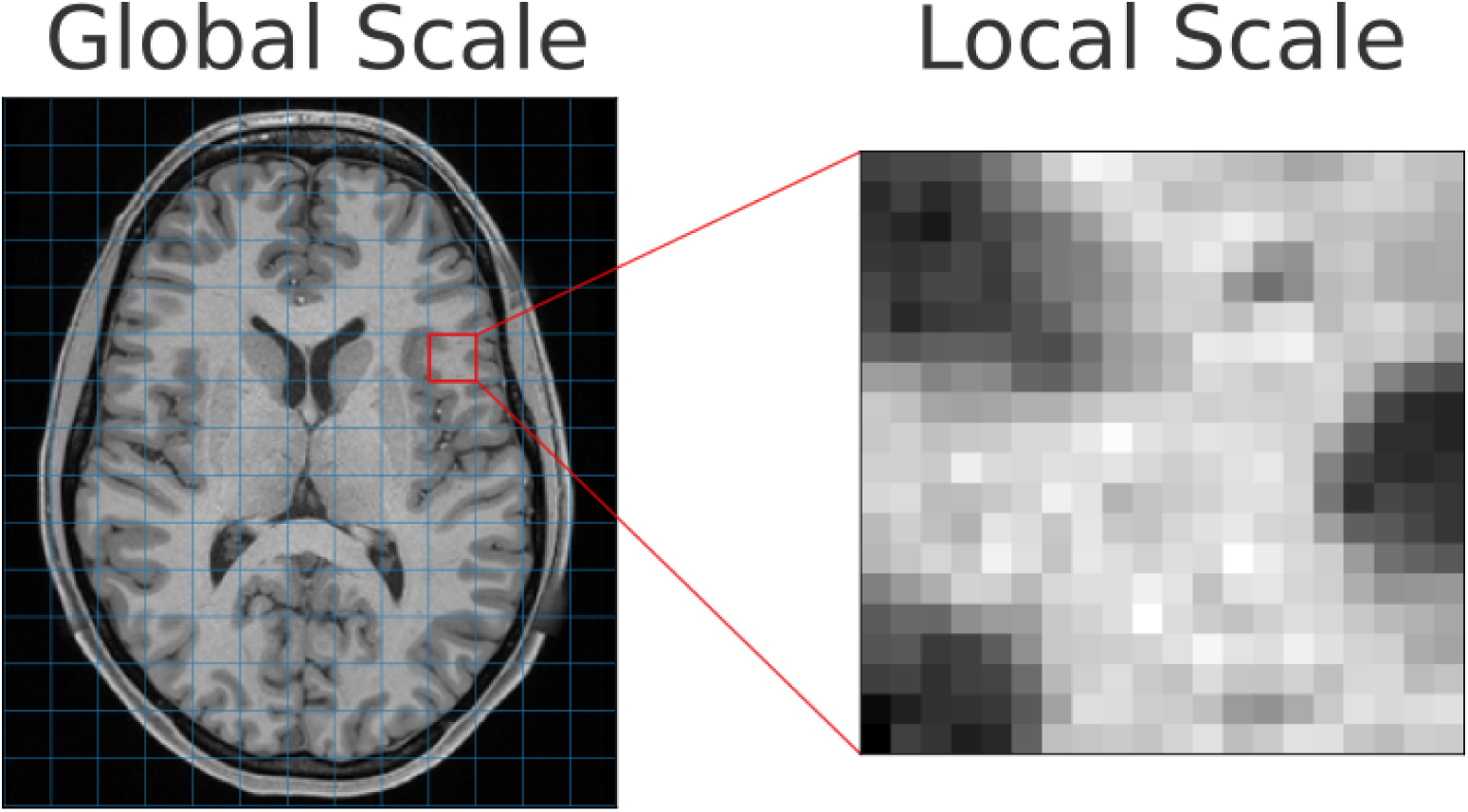
Multi-scale learning. Global features are learned over larger receptive fields (left, blue patches). A convolution neural network (CNN) first compresses the image into a latent feature space. Early layers in the CNN learn local features (right). The feature space is compressed into increasingly global features. The transformer then operates on feature patches (left, blue), with pairwise attention between all patches. These global features embed knowledge related to where a region is with respect to all other regions in the brain; in contrast, local features learn fine boundaries between structures. Multi-scale representations are concatenated using skip connections.

ScaleSurfer includes a second prediction stage that maps anatomical images and predicted labels directly to morphometric features. These include: global measures such as intracranial volume as well as region-wise estimates of volume, cortical thickness, surface area, number of vertices, curvature, and the folding index (Tab. S4). Rather than requiring hours of compute required for surface reconstruction, these models provided estimates in less than a second. This model reuses the same convolutional encoder and transformer bottleneck as the segmentation and parcellation model, replacing the voxel-wise decoder with tabular prediction heads. Multi-scale encoder features are pooled within predicted anatomical labels and concatenated with the global transformer representation. This allows local boundary-sensitive features, regional label information, and whole-brain context to all contribute to each morphometric estimate. This design is motivated by the frequent use of FreeSurfer-derived regional features in machine-learning pipelines, including disease-versus-control classification in Alzheimer’s disease and other disorders.

## 2 Results

### 2.1 Segmentation & Parcellation

Qualitative examples of ScaleSurfer predictions showed close visual agreement with the corresponding FreeSurfer segmentation and parcellation targets (Fig. 3). Dice scores were computed between ScaleSurfer predictions and FreeSurfer (pseudo) ground truth. Representative cases were selected from the test set based on average region-wise Dice scores, corresponding to high, median, and low performance examples near the 95th, 50th, and 5th percentiles, respectively. Even in the lower-performing case, predicted anatomical boundaries remained broadly aligned with the FreeSurfer target labels.

**Fig. 3.**
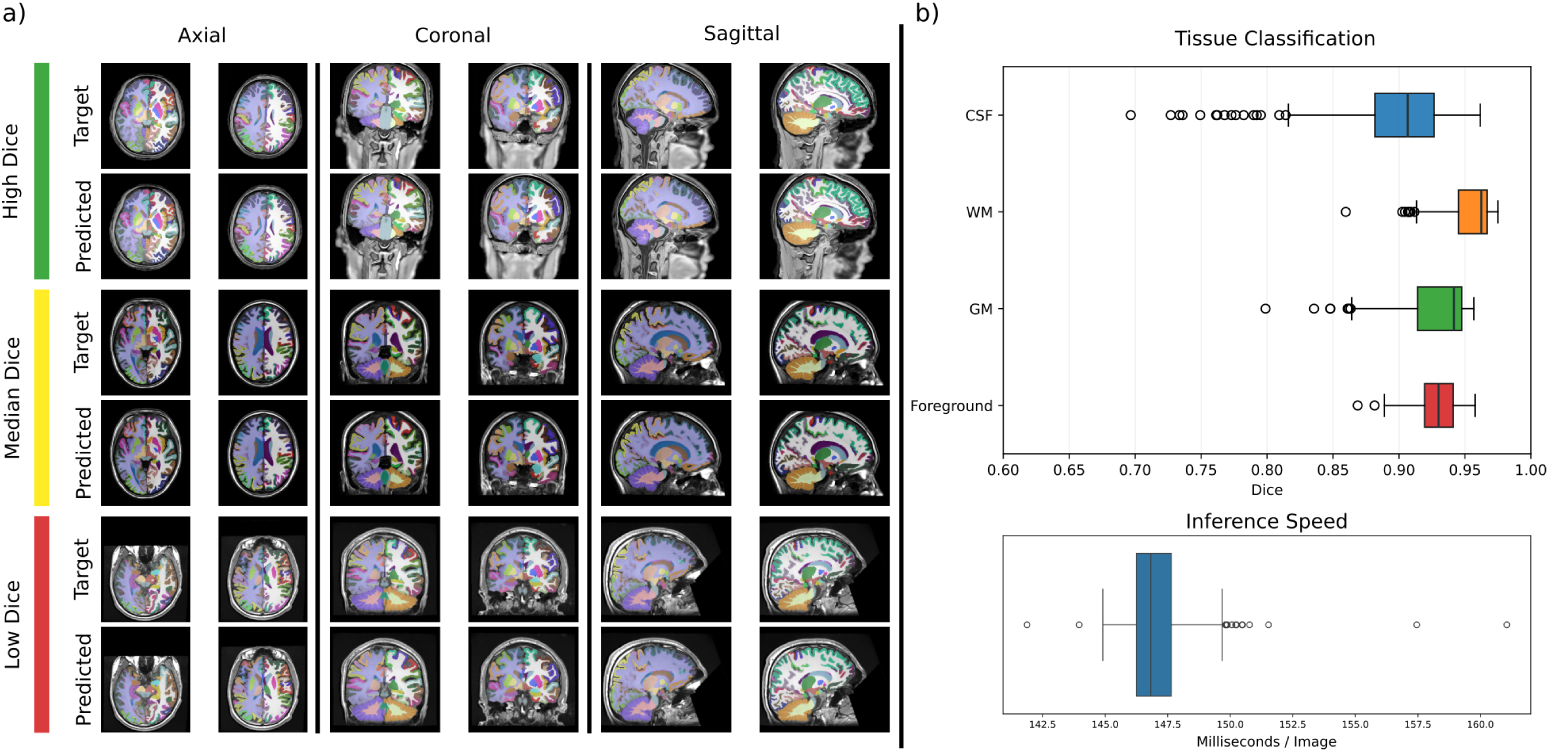
Parcellation and segmentation predictions. **a)** Qualitative test set results show strong alignment between FreeSurfer and ScaleSurfer’s predictions. High dice, median dice, and low dice were defined as the average region-wise dice, closest in percentile to: 0.95 (dice=0.85), 0.5 (dice=0.82), and 0.05 (dice=0.76). Each target-prediction pair (paired rows) corresponds to one image. **b)** Quantitative test set results include dice score and inference speed. FreeSurfer parcellation and segmentation labels were grouped into tissue classes: white matter (WM), gray matter (GM), and cerebral spinal fluid (CSF). These labels were combined into the foreground label. Dice scores were high (mean dice *>* 0.9) for all classes. The mean inference speed was 147 milliseconds per image on a GPU.

Quantitatively, ScaleSurfer achieved high Dice scores across all major tissue classes after grouping FreeSurfer segmentation and parcellation labels into white matter, gray matter, cerebrospinal fluid, and foreground classes (Fig. 3b). Mean Dice scores exceeded 0.9 for all grouped classes, indicating a very strong agreement between the predicted and target anatomical labels. Segmentation and parcellation inference was also fast, requiring an average of 147 milliseconds per image on a GPU when images were processed individually rather than in batches. This represents a speedup of nearly five orders of magnitude over conventional FreeSurfer-style processing, making whole-brain neuroanatomical segmentation and parcellation effectively real time.

That is, ScaleSurfer enables neuroanatomical estimates to be generated immediately after image acquisition.

### 2.2 Morphometric Features

Test set results compare ScaleSurfer predicted features to FreeSurfer-based features, assuming that FreeSurfer segmentations provide some degree of ground truth against which to compare (Fig. 4a). Predicted features are highly correlated with FreeSurfer, at *r >* 0.9 for volume, surface area, thickness, and global measures. Additional surface-based shape measures were more modest in correlation, at *r >* 0.8 for curvature and the folding index. These results show that ScaleSurfer feature predictions are generally correlated with the FreeSurfer (pseudo) ground truth targets.

**Fig. 4.**
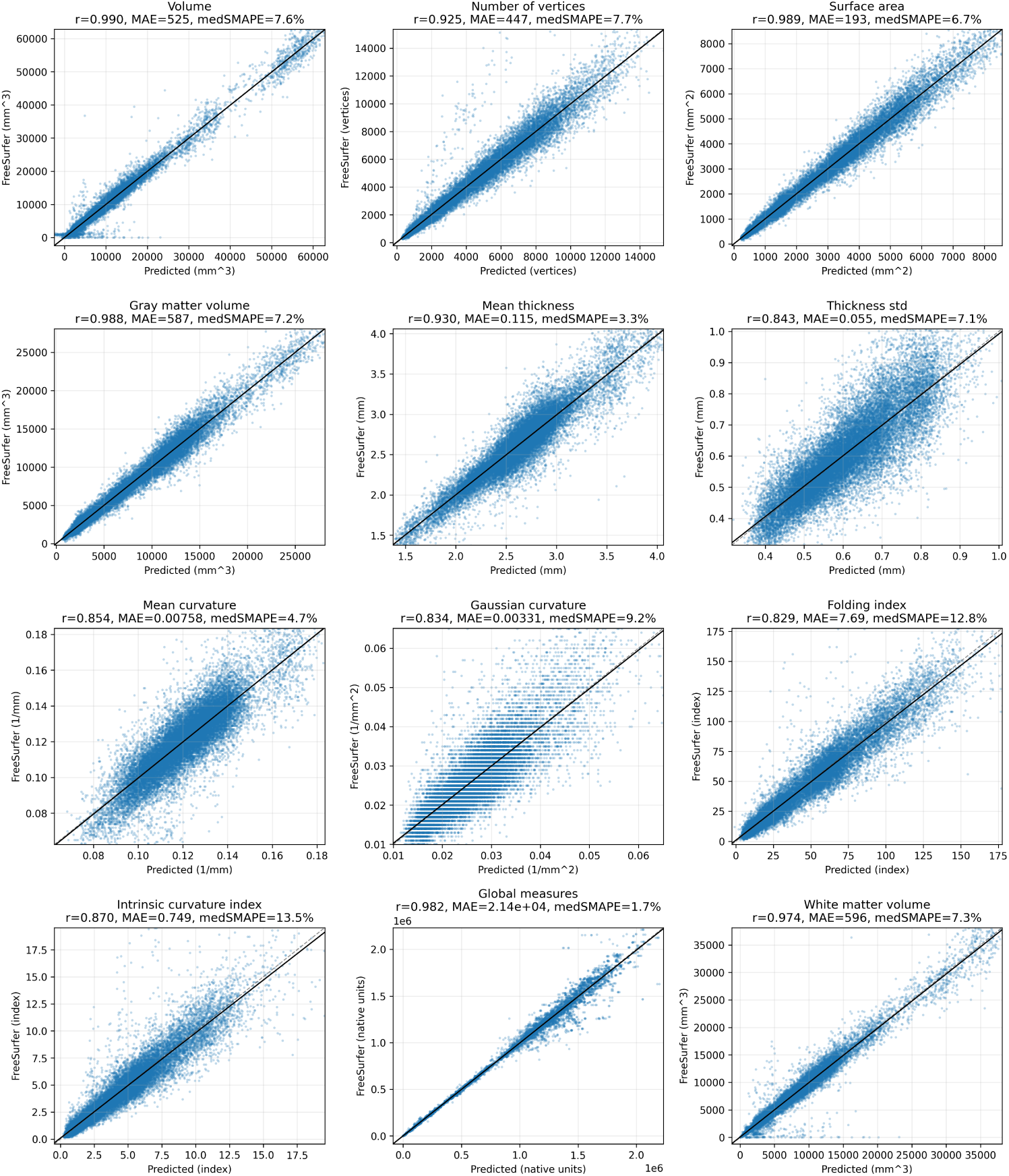
ScaleSufer versus FreeSurfer. Comparison between ScaleSurfer and FreeSurfer metrics on the test set (*n* = 363 subjects, each with 839 morphometric feature predictions). Each point (blue) is a single region-wise feature or global measure. Correlation coefficients ranged between 0.829 and 0.990. Median symmetric mean absolute percentage error (medSMAPE) ranged from 4.7% to 12.8% error. These predicted feature sets are subsequently used to train classifiers.

### 2.3 Clinical Application

In order to assess the utility of ScaleSurfer to real-world applications, we trained a classifier to predict Alzheimer’s disease diagnosis from ScaleSurfer-derived neuroanatomical features. To do this, we used the publicly available Alzheimer’s Disease Neuroimaging Initiative (ADNI) dataset, including 406 patients with an Alzheimer’s disease diagnosis and 1029 healthy controls. Only baseline visit images were included. Notably, the entire dataset was processed – from raw anatomical image, to segmentation and parcellation labels, and finally to feature statistic tables – in 17 minutes (compared to order of weeks needed using more traditional pipelines).

We used a sparse, *L*1-penalized logistic regression model with 5-fold cross validation, and report out-of-fold performance is reported. We assessed performance using the area under the receiver operating characteristic curve (AUROC), which compares the true positive rate – a predicted Alzheimer’s disease diagnosis from ScaleSurfer features when the brain of the scanned person actually has Alzheimer’s disease – against the false positive rate: a predicted Alzheimer’s disease diagnosis when the brain scan was from a healthy control. We observed an AUROC of 0.943, which strongly suggests that ScaleSurfer-derived features capture neuroanatomical variation relevant to Alzheimer’s disease diagnosis (Fig. 5a). When features were residualized, removing the impact of sex, age, and intracranial volume, the AUROC reduced to 0.911. Many of the top-20 ScaleSurfer-derived features include reductions in bilateral medial temporal lobe regions known to be impacted by Alzheimer’s disease (Fig. 5b) [49–54]. These results demonstrate the potential clinical utility of ScaleSurfer-derived features at an incredibly large scale.

**Fig. 5.**
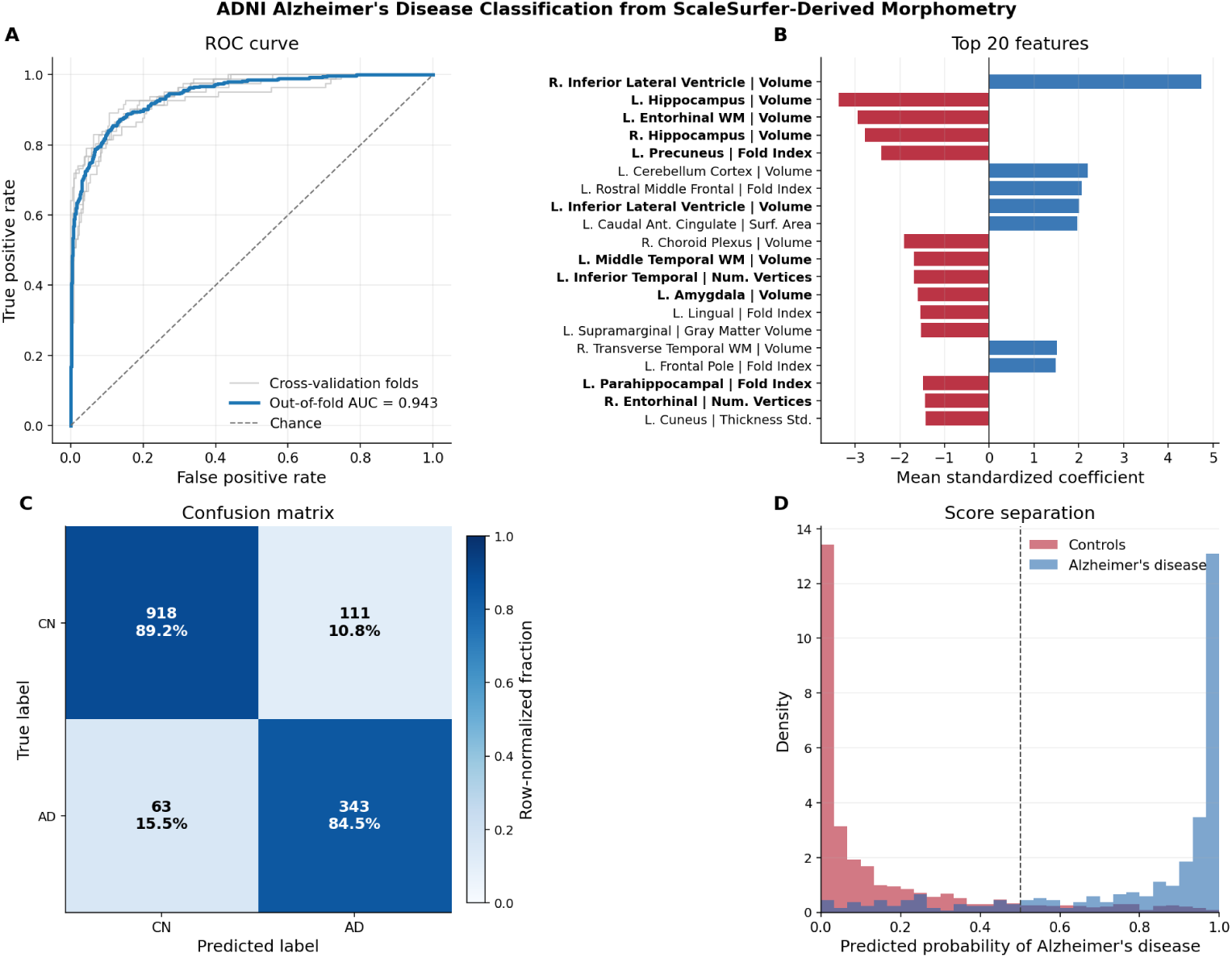
Alzheimer’s disease (AD) classification using ScaleSurfer-derived features. **a)** The receiver operating characteristic (ROC) curve of a sparse logistic regression model trained on ScaleSurfer features, predicting whether or not someone has an Alzheimer’s disease diagnosis based on ScaleSurfer-derived neuroanatomical features. Out-of-fold area under the curve (AUC) was 0.943, demonstrating high discriminative performance in this dataset. **b)** Feature importance based on magnitude of weights. Features in bold are aligned with prior AD literature. **c)** The confusion matrix, raw counts of true positive, false positive, true negatives and false negatives. **d)** Distribution of predicted probabilities of each class.

## 3 Discussion

### 3.1 Importance

In this work, we introduce ScaleSurfer, a fast multi-scale model for structural MRI segmentation, parcellation, and morphometric feature prediction. We demonstrate that a 3D convolutional-transformer architecture can approximate FreeSurfer-derived anatomical labels and regional statistics, but with sub-second processing times that are five orders of magnitude faster than FreeSurfer. We show that ScaleSurfer predicted whole-brain segmentation and parcellation labels have high agreement with FreeSurfer (pseudo) ground truth labels, and that those predicted labels and learned image representations can be used to directly estimate region-wise morphometric features. Finally, we show that ScaleSurfer-derived features support interpretable Alzheimer’s disease classification on an ADNI dataset of 1435 whole brain structural scans – with an end-to-end processing time of just 17 minutes on a single GPU.

These results suggest that detailed structural MRI processing can be reframed as a learned multi-scale prediction problem rather than a sequence of computationally expensive reconstruction and measurement steps. A core motivation for the ScaleSurfer architecture is that anatomical labeling is intrinsically multi-scale, where local image contrast and tissue boundaries are necessary for accurate labeling and broad anatomical context is necessary for learning from global brain structure. ScaleSurfer makes this local / global division explicit, where convolution layers model the local level, and the receptive field grows at each subsequent layer. The transformer models the global level, allowing distant embeddings to interact directly.

In addition to segmentation and parcellation, ScaleSurfer also predicts morphometric features for each region. The decoder is replaced with small model heads that predict regional volumes, cortical thickness, surface area, number of vertices, curvature, and the folding index, as well as global measures.

### 3.2 Limitations

One major limitation is that ground truth segmentation, parcelation, and labels are not really known; instead we rely on FreeSurfer outputs as (pseudo) ground truth targets. So the reported Dice scores and morphometric correlations measure agreement with FreeSurer, rather than absolute anatomical correctness. As such, we cannot adjudicate whether disagreements between ScaleSurfer and FreeSurfer reflect an error in one or the other. This is especially important for unusual anatomy, poor image quality, lesions, and atrophy.

There is also heterogeneity and incompatibility across FreeSurfer versions [55], leading to variable supervision targets when training ScaleSurfer. A base ScaleSurfer model was first pre-trained on all pre-computed FreeSufer versions, e.g., outputs provided from OpenNeuro and HCP, since re-computing FreeSurfer across *n* = 4747 images was computationally prohibitive. Version specific models were then trained by fine-tuning the base model on a FreeSurfer version specific targets. Users should therefore interpret each model as approximating a specified FreeSurfer version or target distribution.

Morphometric predictions also have constraints. ScaleSurfer directly predicts FreeSurfer regional and global statistics and does not explicitly reconstruct the surface. Surface based features such as thickness and curvature are approximations rather than directly computed from surface reconstructions. This distinction matters for applications that require vertex-wise maps, surface topology, surface-based registration, or detailed cortical geometry. For group comparisons or tabular machine-learning models, direct prediction of features was shown to be sufficient.

While the Alzheimer’s disease classification experiment demonstrated the feasibility of using ScaleSurfer features in a clinical context, it does not rule out the possibility of confounding structure: scanner manufacturer, cohort composition, class balance, and demographics. Residualizing out age, sex, and intracranial volume reduced performance from AUC=0.943 to 0.911, indicating that the predictive signal is at least partially associated with non-Alzheimer’s disease specific structure. Broader validation across independent cohorts, scanner sites, disease stages, and clinical labels will be required before clinical claims can be made.

Finally, ScaleSurfer and FreeSurfer are designed around 1 mm isotropic voxels. This choice maximized available training data and made computation more tractable. However, this voxel size limits the ability to learn from higher-resolution anatomical image acquisition. Given the massive speed up that ScaleSurfer provides, future work should explore ways to model and segment the brain at finer resolutions.

### 3.3 Future

Broader validation will help ScaleSurfer to better generalize to additional datasets and populations. Increasing the diversity of training data is aided by open data repositories such as OpenNeuro. As more data is made publicly accessible, it is important to periodically update and improve models. A another clear next step is to extend the ScaleSurfer framework from region-wise morphometric prediction to direct surface reconstruction. It is becoming increasingly clear that efficient models can learn cortical surfaces directly [56–61]. Integrating these approaches with ScaleSurfer would make it possible to preserve the speed of model-based inference while producing more anatomically faithful estimates of surface-derived features.

Current approaches, even deep learning enabled adaptions to FreeSurfer and similar methods, still impose significant time constraints on data processing. These constraints can result in substantial data loss, because poorer quality data – which is especially prevalent among developmental and clinical populations – might not be discovered until well after the scanning session has ended. Current MRI workflows usually measure brain structure in one scan, followed by functional and / or diffusion imaging afterward. The sub-second segmentation enabled by ScaleSurfer paves the way for a totally different approach, where every data sample can be interpreted with an updated estimate of anatomical source, tissue composition, motion, and distortion. That is, if incorporated into MRI data acquisition pipelines, ScaleSurfer could improve image acquisition, preprocessing, and quality control.

Finally, ScaleSurfer makes structural MRI analysis practical at the scale of modern neuroimaging. Long processing times impose significant barriers to scale, creating a bottleneck at a time when it’s becoming more clear that brain-wide association studies leveraging (f)MRI might require thousands of participants worth of data [62]. Fast inference means that large public datasets can be processed in minutes rather than months, turning feature extraction from a bottleneck into an iterative step in model development and analysis. The same framework can be extended to new atlases, alternative parcellation schemes, and harmonized morphometric targets. More generally, ScaleSurfer suggests that fast structural MRI analysis can be built around learned multi-scale representations that produce both anatomical labels and the regional measurements most often used downstream.

## 4 Methods

### 4.1 Data

Training data was collected from OpenNeuro (n=3874) [63] and the Human Connectome Project (n=873) [64]. This provided 4747 total images across 536 studies. Data were organized as paired image and label tensors at the subject level and 80/10/10 splits were used for training, validation, and testing. FreeSurfer parcels and segments (aparc+aseg.mgz) were precomputed by the authors of each dataset. The aparc+aseg.mgz target was chosen since it is default in FreeSurfer and maximized the training set size. This image contains 118 class labels and spans the entire brain, including cortical, sub-cortical, and white-matter regions.

The parcellation and segmentation target images spanned FreeSurfer versions 5 (n=900), 6 (n=1996), 7 (n=1369), and 8 (n=482). Each image in the version 8 set came from a unique OpenNeuro study, promoting diversity in study populations, contexts, scanner manufacturers, and acquisition dates. Version 8 images were processed with FreeSurfer from. All other FreeSurfer outputs were directly fetched from OpenNeuro. Recomputing the entire dataset with one FreeSurfer version would take approximately 800 days assuming four hours per image. ScaleSurfer takes approximately 150 milliseconds per image, or 12 minutes total for the entire dataset, to predict parcellation and segmentation labels.

A base model was trained on the initial set, including segmentation targets from version 5, 6, and 7. The compatibility between versions 6 and 7 has been found to be high, however version 5 is less compatible [55]. FreeSurfer version 8 is also not closely compatible to past versions, for example, the CSF label in version 8 was extended to include the subarachnoid space. To account for version specific nuance, the base ScaleSurfer model was fine-tuned on each version. This resulted in four models additional models, specific to FreeSurfer versions 5, 6, 7, and 8.

For statistics prediction, the supervised target set was restricted to subjects with available FreeSurfer stats directories. The required files were aseg.stats, lh.aparc.stats, and rh.aparc.stats; optional files, including wmparc.stats, brainvol.stats, and aparc+aseg.stats, were parsed when present. Each text file was converted into a canonical long-format table with fields for subject, source file, anatomical region, measure, and numeric value, then pivoted into a fixed target matrix. Supported targets included volumetric measures from aseg and wmparc, cortical regional measures from aparc, and global FreeSurfer summary measures (Tab. S4). This yielded 3,624 supervised statistics samples with an 80/10/10 train/validation/test split of 2,897/364/363 subjects. Version-specific counts were 1,995 for version 6, 1,120 for version 7, and 482 for version 8.

The Alzheimer’s Disease Neuroimaging Initiative (ADNI) dataset was used as an external downstream case study rather than as training data for either the segmentation model or the statistics prediction model. The ADNI metadata contained 1,523 T1-weighted MRI entries, including 1,029 cognitively normal controls, 406 Alzheimer’s disease cases, 78 subjective memory concern cases, 8 mild cognitive impairment cases, and 2 early mild cognitive impairment cases. The primary binary classification analysis used the cognitively normal and Alzheimer’s disease groups only. ScaleSurfer-derived statistics were extracted for each subject and used as tabular morphometric features to evaluate whether the predicted FreeSurfer-like measurements retained clinically relevant anatomical signal.

### 4.2 Model

ScaleSurfer is a three-dimensional encoder-decoder with a 3D-TransUNet-like high-level structure. A convolution hierarchy extracts local anatomical features across spatial scales, a transformer bottleneck integrates long-range context in a compressed latent space, and a decoder reconstructs dense whole-brain label predictions [28, 44, 48]. Exact layer dimensions are reported in Tab. S1. Model implementation and analyses are publicly available at: https://github.com/voytekresearch/scalesurfer

For statistics prediction, the pretrained ScaleSurfer segmentation model was reused as a frozen anatomical feature extractor. The dense segmentation decoder was not used directly for scalar prediction. Instead, an encoder adapter exposed multi-scale feature maps from the convolutional hierarchy and transformer bottleneck. The transformer bottleneck was also globally averaged to provide a whole-brain latent vector, aiding prediction of global features. Predicted parcellation and segmentation labels were used to perform region-aware pooling. For each encoder feature map, the segmentation was nearest-neighbor resampled to the corresponding spatial resolution, and feature activations were grouped by anatomical label. Within each label, channel-wise mean, sum, and standard deviation were computed. Mean pooling summarizes local tissue representation, sum pooling preserves label-size information, and standard deviation captures within-region heterogeneity. These pooled representations were concatenated across labels, scales, and pooling statistics, then concatenated with the global transformer representation.

The statistics model used grouped output heads rather than one independent model per scalar target. A shared pooled feature vector was passed to four task heads: a volumetric head for aseg/wmparc-like regional volumes, a left-hemisphere aparc head, a right-hemisphere aparc head, and a global-measures head. Each head consisted of layer normalization, a fully connected hidden layer with 512 units, GELU activation, dropout of 0.1, and a final linear layer matching the number of targets in that group. This design preserves shared anatomical representation while allowing each target family to learn a specialized mapping from image-derived regional features to FreeSurfer-like scalar outputs.

### 4.3 Training

Preprocessing was intentionally lightweight to increase the speed of end-to-end segmentation. Raw anatomical images were only mean and variance normalized. FreeSurfer’s look-up table (LUT) labels were remapped to a dense class space. Preprocessed outputs were cached before optimization.

Statistics prediction targets were normalized using the training-set mean and standard deviation for each scalar target. Missing targets were masked and did not contribute to the loss. The model was trained with Huber loss on normalized targets, using group-mean reduction so that large target families did not dominate optimization solely by having more scalar outputs. The segmentation encoder was frozen during statistics training, so optimization updated only the lightweight prediction heads.

GPU memory (VRAM) usage was a primary constraint during training. A single, padded 1 mm isotropic brain volume at 208 × 240 × 192 contains 9,584,640 voxels. The network predicts class logits for each output patch element across 118 classes, so the output space is on the order of 1 billion values per image. This estimate excludes parameters from the 14 convolution layers, multi-layer and multi-head transformer, intermediate activations, optimizer state, and gradients, which substantially increase peak memory demand.

Mixed precision was enabled on GPU. To help reduce VRAM load, patch-logit chunking was applied in the output head: the transformer output was converted to patch-level representations and the final linear classifier was evaluated over subsets of patch tokens, with chunk-wise cross-entropy accumulation to reduce peak activation memory. Optimization used AdamW with step-wise cosine annealing and linear warmup. Exact optimization settings are reported in Tab. S3.

## Supplementary information

The main text emphasizes the conceptual structure of the method. Extended background, exact implementation details, optimization settings, and evaluation metrics are summarized below.

### 4.4 UNet

The U-Net architecture uses a symmetric encoder-decoder architecture, from which its characteristic “U”-shape emerges. The encoder progressively shrinks the image through successive convolution and pooling layers, which distill local structure into increasingly global context. Conversely, the decoder expands these compressed features through iterative upsampling, reconstructing spatial fidelity to yield dense, voxel-wise predictions. Skip connections directly link encoder layers to their decoder counterparts and preserve fine-grained spatial detail.

### 4.5 Vision Transformers

Vision Transformers (ViTs) draw inspiration from traditional transformer architectures pioneered in natural language processing (NLP). The input image is first partitioned into fixed-size patches, which are flattened and projected into a latent embedding space to form a sequence of token embeddings. These tokens are transformed through successive self-attention layers, modeling pairwise dependencies across patches. As tokenization inherently omits explicit spatial structure, positional encodings are added to each token embedding to retain each patch’s relative arrangement.

### 4.6 TransUNet

TransUNet is a hybrid encoder-decoder model for medical imaging segmentation, bridging the complementary advantages of prior Transformer and U-Net architectures. First, an encoder uses convolutional layers to extract hierarchical feature maps which capture low-level details (e.g., edges and textures) and high-level features (e.g., regions). These feature maps are flattened into a sequence of token embeddings: stacked together and treated such that each spatial “patch” is a slice through all 2D channels.

Afterwards, these token embeddings are fed through a Transformer encoder, where successive self-attention layers gradually integrate global context. The resultant context-rich embeddings are converted back to a 3D spatial feature map. This feature map is passed into a U-Net decoder; progressively upsampled, while being combined with earlier convolutional features via skip connections to recover fine-grained spatial detail. Finally, a convolutional layer yields a dense, pixel-wise segmentation map for precise localization.

### 4.7 Architecture

ScaleSurfer uses a three-dimensional TransUNet-like encoder-decoder architecture (Tab. S1). A convolutional stem and five stride-2 encoding stages progressively reduce spatial resolution from 256^3^ to 8^3^ while increasing the feature dimension from 12 to 96 channels. Each encoding stage applies downsampling followed by two 3×3×3 convolutions with group normalization and GELU activation. The resulting 8×8×8 bottleneck is reshaped into 512 tokens, augmented with positional encodings, and processed by a two-layer, four-head transformer with feed-forward width 384 to incorporate long-range anatomical context. The decoder restores the original spatial resolution through trilinear upsampling, 1 × 1 × 1 channel reduction, concatenation with the corresponding encoder features, and two 3 × 3 × 3 convolutions with group normalization and GELU activation. Finally, the full-resolution 12-channel representation is normalized channel-wise and passed through a shared linear classifier to produce 118-class logits independently for every voxel. During training and chunked inference, dense voxel features are processed in non-overlapping 16 × 16 × 16 voxel groups, reducing memory use without reducing output resolution.

**Table S1.** ScaleSurfer volumetric segmentation architecture.

| Stage | Operation | Output size |
| --- | --- | --- |
| Input | T1-weighted MRI volume | $1 \times 256 \times 256 \times 256$ |
| Stem | Two $3 \times 3 \times 3$ convolutions, each with GroupNorm and GELU | $12 \times 256 \times 256 \times 256$ |
| Encoder | Five stride-2 $3 \times 3 \times 3$ convolutions; each is followed by two $3 \times 3 \times 3$ convolutions with GroupNorm and GELU | $20 \times 128^3$; $32 \times 64^3$; $48 \times 32^3$; $64 \times 16^3$; $96 \times 8^3$ |
| Bottleneck | Two $3 \times 3 \times 3$ convolutions with GroupNorm and GELU | $96 \times 8^3$ |
| Transformer | Flatten $8 \times 8 \times 8$ latent grid to 512 tokens; add positional encoding; apply 2 transformer encoder layers with 4 heads and feed-forward width 384 | $512 \times 96$ |
| Decoder | Five trilinear upsampling stages; each applies a $1 \times 1 \times 1$ convolution, concatenates the matching encoder feature map, then applies two $3 \times 3 \times 3$ convolutions with GroupNorm and GELU | $64 \times 16^3$; $48 \times 32^3$; $32 \times 64^3$; $20 \times 128^3$; $12 \times 256^3$ |
| Classifier | Channel-wise LayerNorm followed by a linear classifier | $118 \times 256 \times 256 \times 256$ voxel logits |
| Training loss layout | Dense voxel features are reshaped into non-overlapping $16 \times 16 \times 16$ voxel groups before computing the classifier loss | $16 \times 16 \times 16$ groups |

The statistics prediction model reuses the pretrained volumetric TransUNet as a frozen feature extractor. Given a T1-weighted image and a predicted aparc+aseg segmentation, the encoder produces multi-scale feature maps and transformer bottleneck features. For each selected feature scale, the segmentation is resized to the feature grid with nearest-neighbor interpolation, and features are pooled separately within each anatomical label. The pooled representation contains the mean, sum, and standard deviation for every label and feature channel, and is concatenated with a global average-pooled bottleneck vector. This subject-level representation is passed to separate MLP heads for volumetric, left cortical, right cortical, and global FreeSurfer-derived targets. Each head applies layer normalization, a hidden linear layer with GELU and dropout, and a final linear projection to the scalar targets in that output group.

**Table S2.** ScaleSurfer statistics prediction head.

| Component | Operation | Size |
| --- | --- | --- |
| Encoder features | Frozen TransUNet features from **enc1**, **enc2**, **enc3**, **enc4**, and **z** | $20 \times 128^3$, $32 \times 64^3$,<br>$48 \times 32^3$, $64 \times 16^3$,<br>$96 \times 8^3$ |
| Regional pooling | Resize **aparc+aseg** labels to each feature grid; compute mean, sum, and standard deviation within each label | $3 \times 118 \times (20 + 32 + 48 + 64 + 96)$ |
| Global feature | Average-pool transformer bottleneck **z** | 96 |
| Subject vector | Concatenate regional pooled features and global feature | $3 \times 118 \times 260 + 96 = 92136$ |
| Prediction heads | Four MLP heads: LayerNorm, Linear, GELU, Dropout, Linear | **aseg**: 119, **lh_aparc**: 306, **rh_aparc**: 306, **global**: 64 |

### 4.8 Training

The volumetric segmentation training configuration is summarized in Tab. S3. Training used the same compact channel schedule as the architecture in Tab. S1, with deterministic sinusoidal positional encodings and no dropout. Volumes were padded to 256^3^ and trained as full volumes so that the transformer operated with global attention over all 512 bottleneck tokens. The training split was generated once using a per-study 80/10/10 train/test/validation splits. Optimization used AdamW with gradient clipping and mixed precision on CUDA. The learning rate followed a per-step cosine schedule with a one-epoch linear warmup.

The statistics prediction targets are summarized in Tab. S4. The target vector contains 795 scalar FreeSurfer-derived values grouped into four output heads: volumetric aseg/wmparc targets, left cortical aparc targets, right cortical aparc targets, and global summary measures. Cortical targets use the standard aparc table measures for each hemisphere, whereas volumetric targets use Volume mm3 tables. During statistics-model training, scalar targets are normalized using training-set means and standard deviations, and missing targets are masked so that they do not contribute to the loss.

**Table S3.** Volumetric segmentation training configuration.

| Parameter | Value |
| --- | --- |
| Target label set | Default FreeSurfer **aparc+aseg** labels mapped to 118 dense classes |
| Input size after padding | $256 \times 256 \times 256$ |
| Patch size | (16, 16, 16) |
| Channel widths | (12, 20, 32, 48, 64, 96) |
| Transformer depth | 2 |
| Attention heads | 4 |
| Dropout | 0.0 |
| Positional encoding | Deterministic 3D sinusoidal absolute encoding |
| Maximum epochs | 25 |
| Effective batch size | 1 |
| Patch-logit chunk size | 192 |
| Spatial windowing | Disabled; full-volume training |
| Global-attention enforcement | Enabled |
| Optimizer | AdamW |
| Learning rate | $1 \times 10^{-3}$ |
| Minimum learning rate | $1 \times 10^{-6}$ |
| Weight decay | $5 \times 10^{-5}$ |
| Gradient clip norm | 1.0 |
| Learning-rate schedule | Per-step cosine decay with linear warmup |
| Warmup steps | 3388 steps, corresponding to 1 epoch |
| Cosine horizon | 84700 steps, corresponding to 25 epochs |
| Early stopping patience | 6 epochs |
| Target maximum VRAM | 24 GB |
| Mixed precision | Enabled on CUDA |
| Data split | Per-study 80/10/10 manifest; explicit train/validation/test lists |
| Training/validation/test samples | 3388 / 426 / 426 after preprocessing |

**Table S4.**
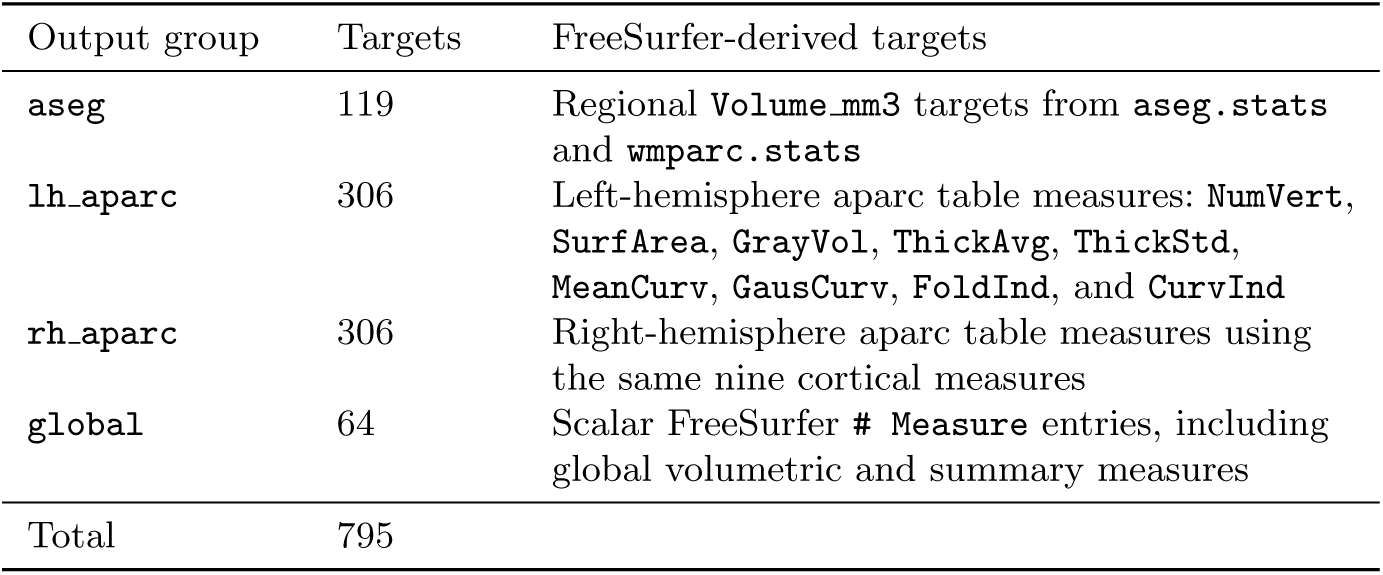
ScaleSurfer statistics prediction targets by output group.

